# Proteomic analysis defines the interactome of telomerase in the protozoan parasite, *Trypanosoma brucei*

**DOI:** 10.1101/2022.11.27.518122

**Authors:** Justin A. Davis, Andres V. Reyes, Nitika, Arpita Saha, Donald J. Wolfgeher, Shou-Ling Xu, Andrew W. Truman, Bibo Li, Kausik Chakrabarti

**Affiliations:** Department of Biological Sciences, University of North Carolina, Charlotte, North Carolina, USA; Department of Plant Biology and Carnegie Mass Spectrometry Facility, Carnegie Institution for Science, Stanford, CA, USA; Department of Molecular Genetics and Cell Biology, The University of Chicago, Chicago IL, USA; Center for Gene Regulation in Health and Disease, Department of Biological, Geological, and Environmental Sciences, College of Arts and Sciences, Cleveland State University, OH, USA

**Keywords:** Telomerase, Telomerase RNA, *T. brucei*, Parasite, proteomics

## Abstract

Telomerase is a ribonucleoprotein enzyme responsible for maintaining the telomeric end of the chromosome. The telomerase enzyme requires two main components to function: the telomerase reverse transcriptase (TERT) and the telomerase RNA (TR), which provides the template for telomeric DNA synthesis. TR is a long noncoding RNA, which forms the basis of a large structural scaffold upon which many accessory proteins can bind and form the complete telomerase holoenzyme. These accessory protein interactions are required for telomerase activity and regulation inside cells. The interacting partners of TERT have been well studied in yeast, human, and *Tetrahymena* models, but not in lower eukaryotes, including clinically relevant human parasites. Here, using the protozoan parasite, *Trypanosoma brucei* (*T. brucei*) as a model, we have identified the interactome of *T. brucei* TERT (*Tb*TERT) using a mass spectrometry-based approach. We identified previously known and unknown interacting factors of *Tb*TERT, highlighting unique features of *T. brucei* telomerase biology. These unique interactions with *Tb*TERT, suggest mechanistic differences in telomere maintenance between *T. brucei* and other eukaryotes.

## 1 INTRODUCTION

Telomeres are the nucleoprotein structures found at the ends of eukaryotic chromosomes. Conventional DNA polymerases are unable to fully replicate the ends of linear DNA molecules, which leads to progressive telomere shortening after every cell division (Shay and Wright, 2019). This problem is solved by the ribonucleoprotein enzyme, telomerase. Proper maintenance of the telomeric end is critical for maintaining genome integrity in eukaryotes. The telomerase enzyme has two essential components: the telomerase RNA (TR), which provides the template required for telomeric DNA synthesis (Greider and Blackburn, 1989); and the catalytic protein telomerase reverse transcriptase (TERT) that catalyzes the de novo synthesis of the telomere G-rich strand. The action of telomerase counteracts progressive telomere shortening after every cell division.

TERT and TR are the minimum components required for the telomerase activity *in vitro* (Chen et al., 2000; Collins, 2006). TR can form a large structural scaffold upon which many accessory proteins can bind to and form the complete telomerase holoenzyme *in vivo*. These accessory proteins are required for telomerase activity and regulation inside of cells (Wang et al., 2019). Interacting partners of the TERT protein have been extensively characterized in yeast, human, and *Tetrahymena* systems, but they have not been extensively studied in lower eukaryotes including clinically relevant human parasites, such as *Trypanosoma brucei* (*T. brucei*). *T. brucei* is the parasite that causes African sleeping sickness in humans. During its life cycle, *T. brucei* will shuttle between an insect vector and a mammalian host. During this time, the parasite will differentiate into distinct developmental stages: Bloodstream form (BF) parasites proliferate in the mammalian host, and Procyclic form (PF) parasites proliferate in the insect vector. Unlike human somatic cells, which tightly regulate telomerase activity, *T. brucei* telomerase is constantly active, enabling continued cell division in their hosts, leading to a chronic infection. BF *T. brucei* also has the advantage of evading its host immune response through antigenic variation. During this process, *T. brucei* regularly switches to express different *Variant Surface Glycoproteins* (*VSGs*), its major surface antigen. *T. brucei* has a very large *VSG* gene pool, but only one *VSG* gene is expressed at any given time. *VSGs* are expressed exclusively from *VSG* expression sites (ESs), which are large polycistronic transcription units located at subtelomeric loci of the parasite’s genome within 2 kb of telomeres (De Lange and Borst, 1982; Hertz-Fowler et al., 2008). Therefore, telomerase activity is critical to maintain the integrity of these *VSG* genes. In cells where the TERT protein has been deleted (*Tb*TERT^-/-^), extremely short telomeres adjacent to the active ES leads to an increase in *VSG* switching frequency (Dreesen and Cross, 2006; Hovel-Miner et al., 2012). Telomeric binding proteins have also been shown to affect *VSG* silencing and switching (Yang et al., 2009; Benmerzouga et al., 2013; Jehi et al., 2014; Jehi et al., 2016; Nanavaty et al., 2017; Afrin et al., 2020; Rabbani et al., 2022). Telomerase mediated telomere maintenance in *T. brucei* is required for the maintenance of subtelomeric *VSG* genes. Because of this, studies of telomerase function and regulation in *T. brucei* could give novel insights into the pathogenicity of this parasite.

The RNA component of *T. brucei* telomerase (*Tb*TR) has a unique structure and sequence composition compared to higher eukaryotes (Sandhu et al., 2013; Podlevsky et al., 2016; Dey et al., 2021). Our recent study on *Tb*TR suggests mechanistic differences in telomere maintenance between *T. brucei* and higher eukaryotes (Dey et al., 2021), underscoring the importance of investigating the functional interactome of *T. brucei* telomerase. Previous proteomic studies on ciliated single-cell protozoan *Tetrahymena* identified several RNA binding proteins, including p65 that copurifies with TR and TERT and promotes proper folding of TR for telomerase holoenzyme assembly and activity (Witkin and Collins, 2004; Berman et al., 2010; Singh et al., 2012; Upton et al., 2017; He et al., 2021). Although homologs of p65 have not been found outside ciliate ancestry, the complex of dyskerin, NHP2, NOP10, and GAR1 that bind the H/ACA domain of human TR are thought to be the functional analog of the p65 chaperone in human telomerase (Berman et al., 2010; Roake and Artandi, 2020). Indeed, these H/ACA binding proteins are known to facilitate human TR folding by enabling the human CR4/5 domain to adopt a particular conformation that interacts with TERT (Egan and Collins, 2012; Chen et al., 2018). Interestingly, in *Tb*TR, the human H/ACA type snoRNP binding domain is replaced by a unique C/D box snoRNA domain (Gupta et al., 2013). In addition, TCAB1, another telomerase RNA-binding protein, was previously discovered as a part of human telomerase complex by mass spectrometry, which is important for intracellular trafficking (Venteicher et al., 2008) and regulating the folding of CR4/5 domain of human TR and telomerase activation. However, despite the fact that *Tb*TR possesses stage-specific structural changes in an active telomerase complex (Dey et al., 2021), major interactors in this complex remain unidentified.

In addition to the canonical function of the TERT protein to protect telomeric ends of chromosomes, emerging evidences also suggest that telomerase can contribute to oxidative stress response in a telomere-independent manner. TERT has been shown to shuttle to mitochondria under increased oxidative stress and influence processes related to DNA damage and cell death (Santos et al., 2004; Ahmed et al., 2008; Indran et al., 2011). There are 5 respiratory chain complexes in human mitochondria that can regulate redox processes and hTERT enhances complex I activity (Ale-Agha et al., 2021). Beyond this, very little is known about the involvement of mitochondrial proteins in TERT function. Interestingly, *T. brucei* protein *Tb*UMSBP2 (Klebanov-Akopyan et al., 2018), which binds to single-stranded G-rich sequence at the replication origins of the mitochondrial DNA of trypanosomatids, colocalizes with telomeres at the nucleus, but whether this activity is coordinated by telomerase mediated DNA repair is not known.

In order to gain a global view and mechanistic insight into telomerase function and regulation in *T. brucei*, we identified the interacting factors of *T. brucei* telomerase reverse transcriptase (*Tb*TERT) using an affinity-purification based mass spectrometry approach. We identified previously known and novel interactors of *Tb*TERT and validated several key interactions. Studying the interactome of *Tb*TERT lays the foundation for future studies of telomerase regulation in *T. brucei*.

## 2. MATERIALS AND METHODS

### 2.1 Culture of Bloodstream Form (BF) *T. brucei* cells

*T. brucei* Lister strain 427 was used throughout this study. All BF cells were grown in HMI-9 media supplemented with 10% heat-inactivated Fetal Bovine Serum (FBS) at 37°C and 5% CO_2_. *T. brucei* Lister 427 strain expressing the T7 polymerase and Tet repressor (single marker, AKA SM) (Wirtz et al., 1999) was grown in media containing 2 μg/ml of G418; *Tb*TERT-FLAG-HA-HA (F2H) cells grown with 2 μg/ml of G418, 0.1 μg/ml Puromycin; *Tb*TR ΔC/D box mutant cells were grown with 2 μg/ml of G418, 4 μg/ml of Hygromycin, 2.5 μg/ml of Phleomycin, 5 μg/ml of Blasticidin, 0.1 μg/ml of Puromycin and 0.1 μg/ml of Doxycycline to constitutively induce the *Tb*TR mutations.

### 2.2 Plasmids

The *Tb*TR WT gene without 3’ C/D box region (nt 841–943) together with 400 bp upstream and 380 bp of downstream *Tb*TR flanking sequences were cloned into the pLew111 plasmid to generate pLew111-*Tb*TR ΔC/D box plasmid

### 2.3 Generation of the BF *T. brucei* ΔC/D box mutant strain

To generate the *Tb*TR ΔC/D box mutant strain, pLew111-*Tb*TR ΔC/D box plasmid was digested with NotI and targeted to an rDNA spacer in the SM/*Tb*TR^-/-^ cells under the phleomycin selection. Clones were confirmed by northern analysis. XhoI digested pSK-*Tb*TERT-3C-FLAG-HA-HA-*PUK* plasmid (Dey et al., 2021) was subsequently transfected into the same cells under the puromycin selection to generate the SM/ *Tb*TR ΔC/D box/ *Tb*TERT ^+/F2H^ strain. Clones were confirmed by western and Southern blotting

### 2.4 Immunopurification of *T. brucei* Telomerase Complexes

Immunoprecipitation of *T. brucei* telomerase was performed using a custom made anti-*Tb*TERT antibody (Dey et al., 2021) to purify native telomerase complexes from BF Wild-type (WT) and BF *Tb*TR ΔC/D box cells. Approximately, 5 X 10^8^ cells/300 ml were collected by centrifugation at 1900 RPM for 6 minutes. Following centrifugation, cells were lysed by homogenization in 500 μl of 1X immunopurified (IP) lysis buffer (25 mM Tris-HCl pH 7.5, 150 mM KCl, 1 mM EDTA, 10 mM MgCl_2_, 0.5% IGEPAL CA630, 1X protease cocktail inhibitor, and 20 units of Ribolock RNase inhibitor). Lysate was then cleared of cell debris by centrifugation at 3000 RPM for 5 minutes at 4°C. The lysate was then pre-cleared by incubated with 50 μl of pre-washed Dynabeads protein G (10003D) for 1 hr. at 4°C on rotation. Pre-cleared lysates were then incubated overnight at 4°C on rotation with 5 μg of a custom anti-*Tb*TERT antibody and an IgG antibody was added to the control (Dey et al., 2021). The next day, 50 μl of pre-washed Dynabeads protein G was added to the lysate antibody mixture and incubated at 4°C for 2 hrs. on rotation. After incubation, the beads were collected in a magnetic stand and washed twice in 1X IP lysis buffer. After washing, the bound protein was eluted off the beads by boiling in 100 μl of 1X SDS-PAGE dye for 5 minutes at 95°C. Eluted proteins were then stored in −80°C until further use. Each experiment was performed in biological triplicate. Bound complexes were assayed for the presence of *Tb*TERT by using an anti-FLAG antibody as the BF WT cells were *Tb*TERT-FLAG-HA-HA tagged. Briefly, 4 μl of 100 μl of sample was loaded onto 4-12% Novex Tris-glycine gel (Invitrogen, XP04120BOX). Western blotting was done using an anti-FLAG antibody, diluted 1:500, and the VeriBlot for IP Detection Reagent (HRP, ab131366) diluted to 1:10000. Detection was then done using Pierce ECL Plus Chemiluminescence kit (Thermo Fisher Scientific, 32106). Imaging was then done using Bio-Rad ChemiDoc MP system.

Immunopurification was also performed using Pierce Anti-DYKDDDDK magnetic beads (A36797). Approximately 6 × 10^8^ cells/300 ml were harvested and lysed in 300 μl of immunopurified (IP) lysis buffer (25 mM Tris-HCl pH 7.5, 150 mM KCl, 25 mM NaCl, 1 mM EDTA, 10 mM MgCl_2_, 0.5% IGEPAL CA630, 1× protease cocktail inhibitor and 20 units of Ribolock RNase inhibitor). Lysate was cleared of debris by centrifugation at 3000 rpm for 5 min at 4°C and incubated with pre-washed 50 μl of Pierce Anti-DYKDDDDK magnetic beads (A36797) at 4°C for 2 h with rotation. Following incubation, the beads were washed twice by ice cold IP buffer and once with ice cold DEPC water. The beads were then resuspended in 50 μl of RNAse free water.

### 2.5 LC-MS/MS analysis

Proteins were separated by SDS-PAGE and Gel segments were cut and subjected to in-gel digestion using trypsin. Peptides were desalted using C18 ZipTips (Millipore). Peptides were analyzed on a Q-Exactive HF hybrid quadrupole-Orbitrap mass spectrometer (Thermo Fisher) equipped with an Easy LC 1200 UPLC liquid chromatography system (Thermo Fisher). Peptides were first trapped using trapping column Acclaim PepMap 100 (75 uM x 2cm, nanoViper 2Pk, C18, 3 μm, 100A), then separated using analytical column Acclaim PepMap RSLC (75um x25cm, nanoViper, C18, 2 μm, 100A) (Thermo Fisher). The flow rate was 300 nL/min, and a 120-min gradient was used. Peptides were eluted by a gradient from 3 to 28% solvent B (80% (v/v) acetonitrile/0.1% (v/v) formic acid) over 100 min and from 28 to 44% solvent B over 20 min, followed by a short wash at 90% solvent B. For DDA acquisition, the precursor scan was from mass-to-charge ratio (m/z) 375 to 1600 and the top 20 most intense multiply charged precursors were selected for fragmentation. Peptides were fragmented with higher-energy collision dissociation (HCD) with normalized collision energy (NCE) 27.

### 2.6 LC-MS/MS data and statistical analysis

The resulting raw data files were searched against a concatenated library (TbruceiTREU927 release 32 databases with 11202 entries) using MaxQuant. Carbamidomethyl Cysteine was set as a fixed modification. Oxidation of methionine and N-terminal acetylation were set as variable modifications. Tolerance for precursor ions was set to 4.5 ppm and 20 ppm for fragment ions. A maximum of two missed cleavages was allowed. MaxQuant was set to match in between runs and report LFQ. All other parameters were at the default setting.

The proteinGroups.txt file generated by MaxQuant was further processed using Perseus. Reverse and possible contaminants were removed from the protein groups. Samples were separated into an experimental group consisting of the pull downs of isotype matched IgG (control) and a group consisting of the TERT Immuno-Precipitation pull downs. Protein groups were filtered to contain at least three quantifications in one experimental group. The remaining missing quantifications were imputed with random numbers from a normal distribution (width 0.3, shift = 1.8). A two-sided student t-test was performed across replicates between each experimental group.

### 2.7 MS bioinformatics analysis

The mass spectrometry proteomic data was analyzed by a range of approaches. Volcano plot was generated using GraphPad Prism software version 9.3.1. The STRING database was used for classifying proteins based on functional categories and gene ontology (GO) terms. Protein-protein interaction network analysis was done using STRING version 11.5 (https://string-db.org/) and visualized by using the Cytoscape software version 3.9.1.

### 2.8 Structure-Guided Predictions

Proteins identified by mass spectrometry contained several hits that have very little primary sequence identity with proteins from other phyla, so their functional orthologs were not entirely evident from simple sequence homology. To identify whether structural homology exists between the local folds of these proteins and those reported in other organisms, we obtained predicted structure models of these *T. brucei* proteins from AlphaFold (Jumper et al., 2021) using their Uniprot IDs and then queried these ‘PDB’ entries using programs PDBeFold (Krissinel and Henrick, 2004) and DaliLite (Holm and Rosenstrï¿ ½m, 2010). The top scoring and only relevant hit with a DaliLite Z score cut-off of >2 is considered as biologically informative structural neighbor of the protein of interest (**Supplementary Table S1**).

### 2.9 Western Blotting and SDS-PAGE analysis

All *Tb*TERT western blots were done using either an anti-FLAG antibody (1:500) or a custom anti-*Tb*TERT C terminus antibody (1:500) unless otherwise indicated. Nucleolar protein 58 (NOP58) was detected using an anti-NOP58 antibody (Thermo Fisher Scientific, PA5-54321) diluted 1:500 and an anti-Rabbit HRP conjugated secondary antibody diluted to 1:10000. To qualitatively check protein levels, 4 μl of IP eluate was separated on a 4-12% Novex Tris-glycine gel and stained with Coomassie Brilliant Blue R-250 Dye (Thermo Scientific, 20278) for 30 minutes. The gel was then destained until bands were visible in destain solution (40% MeOH, 10% acetic acid).

### 3.0 *Tb*TR detection and Telomerase activity assay

*Tb*TERT IP was performed as described earlier. To detect the presence of *Tb*TR in the IPed complex, total RNA was isolated from the protein G magnetic beads using the TRIzol reagent (Thermo Fisher Scientific, 15596026) following the manufacturers protocol. 100 ng of isolated RNA was then used for cDNA synthesis utilizing the SuperScript II reverse transcriptase (Thermo Fisher Scientific, 18064022) following the manufacturers protocol. The generated cDNA was then used for qRT-PCR analysis using *Tb*TR specific primers (Fwd: CTGTGGAAATTTGTCGTAAGTG, Rev: AGTAGGGTTAGGGATCGTATAG).

To determine the activity of the Immunopurified *T. brucei* telomerase complex, a modified version of the exponential isothermal amplification of telomere repeat (EXPIATR) assay was performed (Tian and Weizmann, 2013; Dey et al., 2021)Briefly, A master mix was prepared on ice consisting of Nicking Telomerase Substrate (NTS, GTGCGTGAGAGCTCTTCCAATCCGTCGAGCAGAGTT), Nicking Probe (NP, AGCAGGAAGCGCTCTTCCTGCTCCCTAACCCTAACCC), 1X EXPIATR buffer (30 mM Tris-HCl, pH 8.3, 1.5 mM MgCl_2_, 100 mM KCl, 1 mM EGTA, 0.05% v/v Tween20), 200 μM dNTPs, Bst 2.0 Warm start DNA polymerase (0.96 units) and Nt. BspQ1 NEase (5 units). 17 μl of the master mix was aliquoted to PCR tubes containing, 3 μl of anti-FLAG bead-bound *T. brucei* telomerase, RNase A treated or heat-inactivated telomerase RNP bound beads, telomerase positive control (TPC8, GTGCGTGAGAGCTCTTCCAATCCGTCGAGCAGAGTTAGGGTTAGGGTTAGGGTTAGGGTTAGGGTTAGGGTTAGGGTTAGGG) (0.5 μM) and blank beads as a negative control. Telomerase activity was initiated by initial incubation of tubes at 28°C for 45 min for Nicking telomerase substrate (NTS) extension followed by amplification of resultant telomerase products at 55°C for 30 min. The amplified products were then analyzed on 12% Native PAGE gel by loading 10 μl of the reaction mixture.

## 3. RESULTS

### 3.1 Affinity-purification mass spectrometry (AP-MS) of BF *T. brucei* telomerase reverse transcriptase

Characterizing the global interactors of a protein of interest can be done through affinity-purification mass spectrometry (AP-MS). Identifying the interactome of a protein is key in understanding its function in the cell and how it is regulated. To identify the global protein interactors of *T. brucei* telomerase, we utilized AP-MS to identify the global interactome of *Tb*TERT at the BF stage. We first immunopurified (IP) *Tb*TERT using a custom anti-*Tb*TERT antibody along with its associated proteins and performed LC-MS/MS (**Figure 1A**). For immuno-affinity purification, 500 ul of lysate containing 1 mg of total protein was used per IP sample. *Tb*TERT was pulled down from the lysate using a custom anti-C terminus *Tb*TERT antibody. In addition to verifying the specificity of the custom antibody for binding to *Tb*TERT, the presence of some non-specific cross-reactive bands were also observed. The presence of the immunopurified *Tb*TERT was confirmed using western blotting and SDS-PAGE analysis (**Figure 1B-D**). Immunoblot analysis (anti-FLAG antibody) of the IP fractions from *Tb*TERT-F2H cells showed that *Tb*TERT was enriched in the pulldown products from anti-*Tb*TERT C terminus antibody IPs but not from control groups. The anti-*Tb*TERT C-terminus antibody IP samples were then subjected to SDS-PAGE and in-gel protease digestion, followed by mass spectrometry as described in ‘Materials and Methods’. To further validate the presence of telomerase components in the IP, RNA was extracted from IP beads and subjected to qRT-PCR analysis to confirm the presence of the telomerase RNA in the complex (**Figure 1E**). To determine if the immunopurified telomerase complex was catalytically active, we performed a telomerase activity assay (EXPIATR) using the IPed complex (**Figure 1F**). The result confirmed that the immunopurified *T. brucei* telomerase complex was catalytically active. To independently validate our proteomic screen, another set of affinity enrichment of *Tb*TERT protein using an anti-FLAG antibody was performed in duplicates in *Tb*TERT-F2H cells (two biological replicates) and screened for known *Tb*TERT interactors. Since *Tb*TERT and several other proteins which were previously linked to telomerase were identified in both of the above-mentioned IP and mass spec data sets, it was apparent that the ribonucleoprotein complex identified by this approach is biologically relevant to telomerase function.

**Figure 1:**
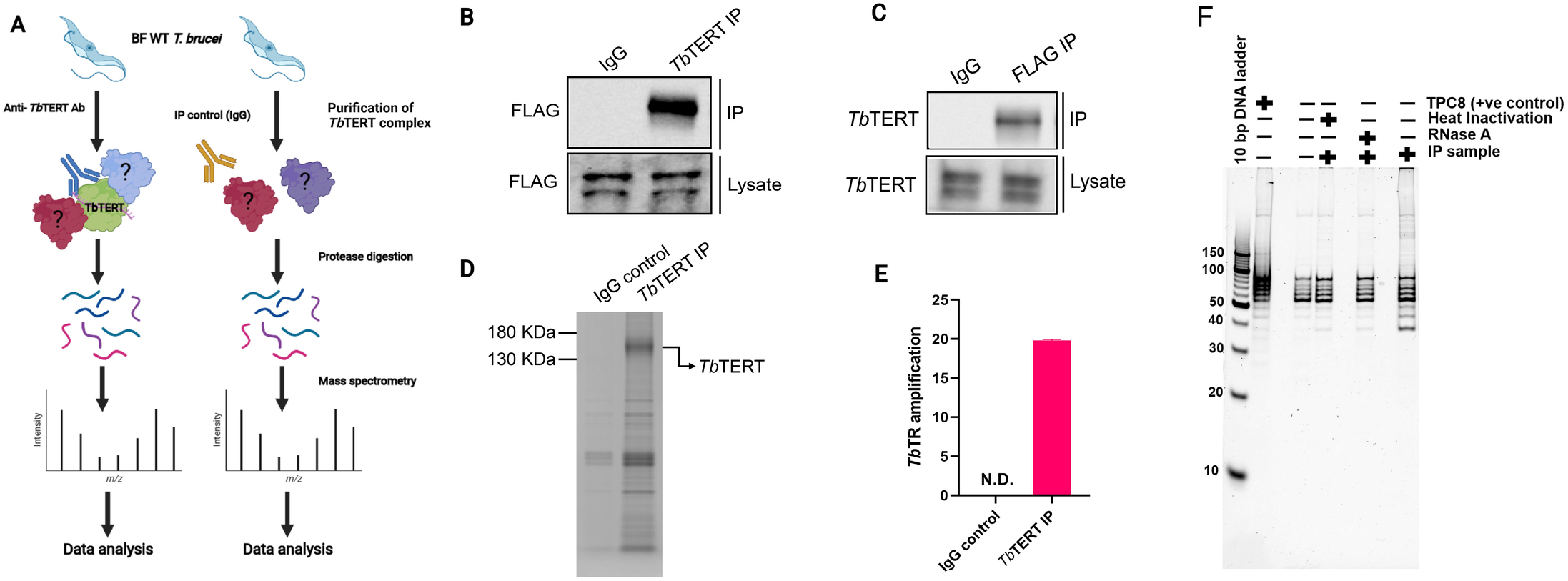
Affinity-purification mass spectrometry of bloodstream form *T. brucei* telomerase reverse transcriptase. (**A**) Experimental workflow for proteomic analysis. BF *T. brucei* cells expressing a FLAG tagged version of *Tb*TERT were grown and *Tb*TERT complexes were purified using a custom anti-*Tb*TERT C terminus antibody. An IgG isotype antibody was used as a control. Purified *Tb*TERT complexes were then digested with trypsin. These peptides were then analyzed by LC-MS/MS. (**B**) Western blot confirming the presence of immunopurified *Tb*TERT. IP samples were obtained and run on SDS-PAGE gels and immunoblotted with anti-FLAG antibody to detect *Tb*TERT. (**C**) Western blot confirming the presence of *Tb*TERT. IP samples were obtained and run on SDS-PAGE gels and immunoblotted with an anti-*Tb*TERT C terminus antibody to detect *Tb*TERT. (**D**) SDS-PAGE analysis of immunopurified *Tb*TERT. A small aliquot was also resolved on SDS-PAGE and stained with Coomassie stain to qualitatively check *Tb*TERT protein levels. (**E**) RT-qPCR detection of *Tb*TR from Immunopurified *Tb*TERT complexes. (**F**) Telomerase activity of the bead-bound telomerase enzyme was analyzed by telomerase primer extension assay.

### 3.2 Proteomic Analysis of BF *T. brucei* telomerase reverse transcriptase

We identified 1056 proteins with 2 or more peptides. After stringent filtering of this dataset for those proteins highly enriched (Log_2_(fold change) > 1.9) in the IP vs control, 66 high-confidence proteins remained. To study the interactome of *Tb*TERT, the relative abundance (log_2_(fold change)) and statistical significance -log_10_(*P*-value) of the proteins from the IP samples and controls were calculated (**Figure 2A**). This resulted in the enrichment of 66 proteins for the anti-*Tb*TERT IP samples. The protein samples that were significantly enriched in the IP samples versus the controls (Student’s *t*-test, -log_10_(*P*-value) >1.3) included both nuclear and mitochondrial proteins, such as *T. brucei* telomerase reverse transcriptase (*Tb*TERT, Tb927.11.10190), yeast telomerase cell cycle turnover-related (anaphase promoting complex) proteins, CDC16 (Tb927.6.2150) and CDC27 (Tb927.10.10330) homologs (Sealey et al., 2011; Ferguson et al., 2013), mitochondrial stress-response protein, human HSP60 homolog (Tb927.11.15040) which is known to accumulate with hTERT in the same fractions of human mitochondria (Sharma et al., 2012), Splicing factor 3B subunit 1 (SF3B1) homolog (Tb927.11.11850) involved in various cellular functions including DNA damage response (Te Raa et al., 2015) and telomere maintenance (Wang et al., 2016), damage specific DNA binding protein 1 (DDB1) homolog (Tb927.6.5110), involved in ubiquitin-mediated TERT protein degradation (Jung et al., 2013), and several other proteins involved in telomerase and telomere metabolism (selected proteins shown in **Figure 2A** and **Table 1**).

**Figure 2:**
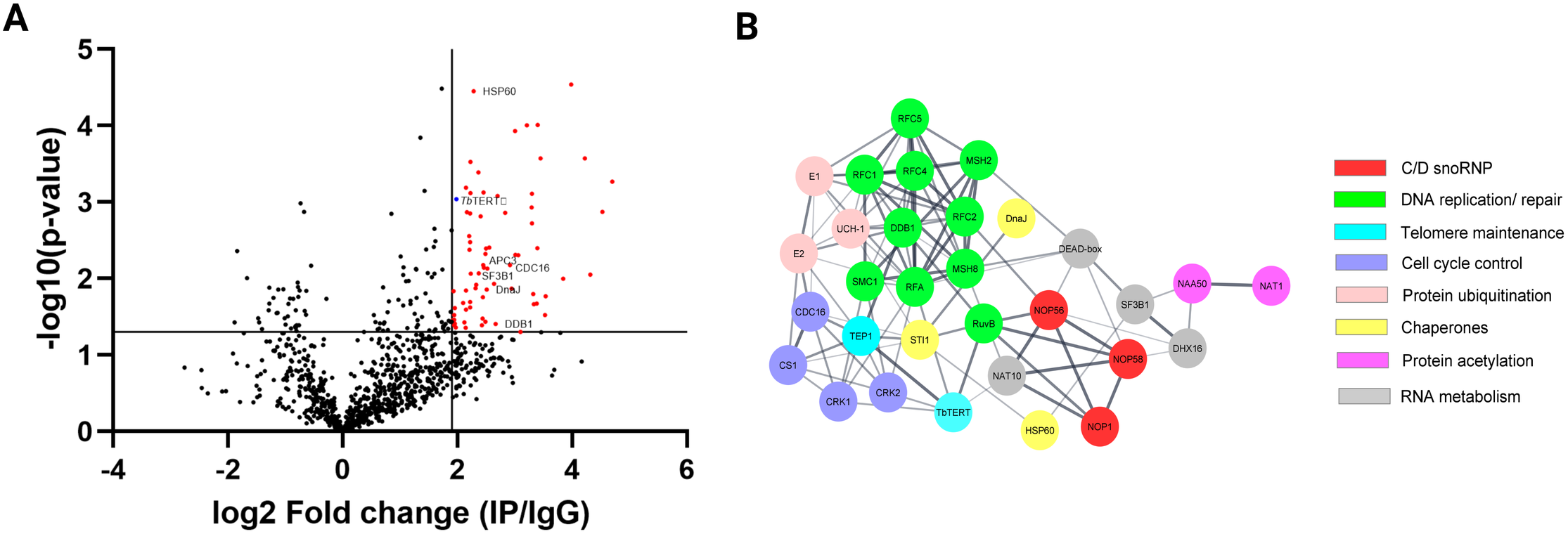
**(A)** Volcano plot was performed with an x-axis representing the difference in logarithmic protein intensities between the *Tb*TERT immunoprecipitation elution and the isotype matched IgG control (Elution and Control experimental groups). The y-axis is the negative log of the two-sided student’s t-test. The volcano plot serves as a visual representation of the protein groups that are significantly enriched between the elution and control groups. These enriched groups contain the bait protein *Tb*TERT and several candidates interacting with a p-value <= 0.05. **(B)** Protein-protein interaction network of relevant *Tb*TERT hits identified by MS. Network was generated using the STRING database and visualized using Cytoscape. Colors of nodes represent the protein’s biological function. The thickness of the lines denotes the strength of the interaction (confidence PPI, threshold: 0.4, medium confidence).

**Table 1.** Significantly enriched proteins identified in *Tb*TERT complex

| Protein | Accession number<br>(UniProtKB) | Spectral counts | Unique peptides |
| --- | --- | --- | --- |
| *Tb*TERT | Q383R0 | 10 | 5 |
| HSP60 | Q381T1 | 6 | 5 |
| DnaJ | Q38C15 | 19 | 5 |
| APC3 | Q389U4 | 21 | 6 |
| CDC16 | Q584U1 | 30 | 12 |
| SF3B1 | Q382Z6 | 26 | 14 |
| DDB1 | Q586I3 | 21 | 13 |

To highlight the connectivity of candidate *Tb*TERT interactors, we used the STRING database and Cytoscape to generate a functional protein-protein interaction network of *Tb*TERT (**Figure 2B** and **Supplementary Figure S1**). To further determine the functions of proteins in the network, STRING GO analysis was done and proteins were grouped by biological process, and cellular component (**Figures 3A-B**). Terms for biological process that were enriched included, telomere maintenance by telomerase, cell cycle control, DNA repair, and response to stress. Enriched cellular component terms included, box C/D snoRNP complex, DNA replication factor C, anaphase-promoting complex, and cullin-RING ubiquitin ligase complex. Notably, proteins that are important for telomerase RNA biogenesis, processing and trafficking were significantly enriched in these GO terms.

**Figure 3:**
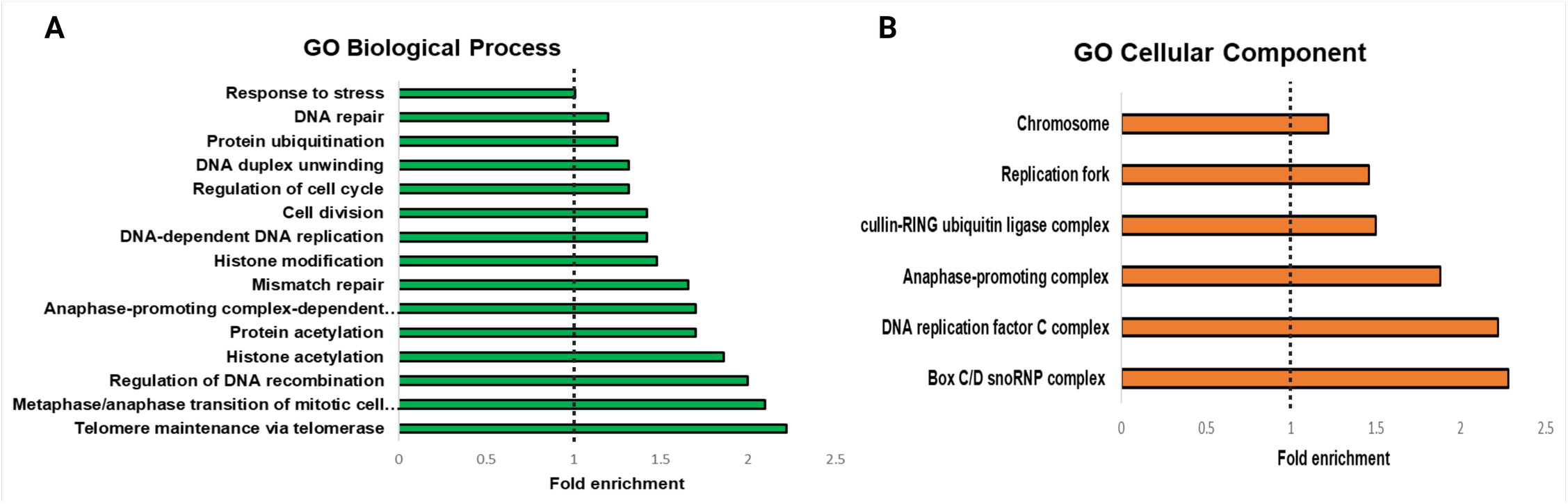
GO analysis of *T. brucei* TERT interactome. STRING GO analysis **(A)** Enrichment by Biological process. The top 15 enriched GO terms are shown. **(B)** enrichment by Cellular component. The top 9 enriched GO terms are shown.

An independent set of IP-MS experiment also validated majority of the proteins identified in the mass spec run described above. A false discovery rate (FDR) of 1% was used as cut off for this data (**Supplementary Table S1**). Both these sets of experiments also identified proteins that have no known relationship with telomerase and therefore these hits could be false positives or newly discovered interactors of *Tb*TERT. For example, an RNA cytidine acetyltransferase, NAT10 homolog (Tb927.5.2530) was found enriched in all proteomic datasets. NAT10 was previously shown to have predominantly nucleolar localization, association with human telomerase, and is primarily involved in telomerase RNA biogenesis (Fu and Collins, 2007). One more example is the putative NOT1 deadenylase (Tb927.10.1510), part of the CCR4-NOT deadenylase complex, which plays important regulatory roles both at the transcriptional and post-transcriptional levels, such as heterochromatic repression of sub-telomeric genes in fission yeast (Cotobal et al., 2015) and rapid deadenylation of m6A-containing RNAs by the CCR4–NOT deadenylase complex in mammalian cells (Du et al., 2016). Given that a majority of expressed virulence genes (*VSGs*) in *T. brucei* are subtelomeric (Saha et al., 2020) and human telomerase RNAs are known to contain m6A signatures (Han et al., 2020), the role of Tb927.10.1510 remains unexplored but relevant to *T. brucei* telomere biology.

### 3.3 Known and novel interactors are part of the active telomerase RNP in *T. brucei*

Homologs of telomerase-associated proteins which are known regulators of telomerase functions were found to be part of the IP telomerase complex in *T. brucei*. Typically, three types of known telomerase-associated proteins were reported to co-purify with TERT. The foremost of the three are the telomerase RNA-binding proteins that are involved in TR biogenesis, trafficking and TR-TERT assembly. These include, for example, dyskerin in vertebrates (Mitchell and Collins, 2000; Mochizuki et al., 2004), Sm proteins in yeasts (Tang et al., 2012), and La-motif proteins, such as p65, in ciliates (Singh et al., 2012). Interestingly, dyskerin binding H/ACA domain of human TR is replaced by a novel C/D box domain in *T. brucei* (Gupta et al., 2013). Several unique C/D box snoRNA binding proteins (snoRNPs) were identified in our *Tb*TERT immunopurified complex which are described in the next section. In addition, a *T. brucei* La protein, Tb927.10.2370, which shows 24% amino acid sequence identity and a Z-score of 8.6 with *Tetrahymena thermophila* TR binding protein p65 was also identified in the IP-MS. Telomerase RNP assembly also requires molecular chaperones, such as AAA+ family of ATPases, known as Pontin and Reptin, which can directly interact with TERT and play critical roles in telomerase RNP accumulation (Venteicher et al., 2008). Both the Pontin (Tb927.4.1270) and Reptin (Tb927.4.2000) homologs, annotated as RuvB-like DNA helicases in the *T. brucei* genome database, were also identified by this *Tb*TERT AP-MS analysis. Additionally, another AAA+ ATPase protein, a yeast CDC48 homolog, Tb927.10.5770, was also identified. CDC48, which was previously identified in a molecular complex that recognized and bound ubiquitinated proteins (Schuberth et al., 2004), was also found to be associated with yeast telomerase as a novel regulator of telomere length homeostasis. Notably, TERT turnover is dependent on ubiquitin-proteasome mediated degradation process (Jung et al., 2013) and therefore proteins that are important for ubiquitination were previously identified in the telomerase complexes, such as several isoforms of E3 ubiquitin ligases (Lin et al., 2015). Interestingly, several ubiquitin-family proteins were also identified in the IP-MS data including ubiquitin ligases, although the roles of these proteins in *T. brucei* telomerase biology remains uncertain until the ubiquitination status of *Tb*TERT is determined. Mammalian studies have identified chaperone proteins p23 and HSP90 as two important proteins that are physically and functionally associated with telomerase activity (Holt et al., 1999). The proteomic mapping also identified a mammalian HSP90 homolog, Tb927.3.3580, with enrichment of several unique peptides in *Tb*TERT IP samples identified by MS. Our proteomics screen also identified a *T. brucei* homolog of STI1 (Tb927.5.2940), which is a major co-chaperone of HSP90. Poly(A)-specific ribonuclease (PARN) is a 3’-exoribonuclease that is known to play important role in the maturation of telomerase RNA (Moon et al., 2015). A *T. brucei* homolog of PARN, Tb927.9.13510 was identified in this proteomic mapping data that may relate with the fact that *T. brucei* telomerase RNA is a Pol II transcript (Sandhu et al., 2013) that may require PARN processing for maturation. All these known telomerase homologs of *T. brucei* are listed in Supplementary Table – S1. Intriguingly, 15 different nucleoporins were also identified in this proteomic screen which may promote telomeric chromatin organization or nuclear import of *Tb*TERT (Frohnert et al., 2014; Pinzaru et al., 2020). In terms of proteomic identification, it should be noted that several of the above proteins were identified in the range of low scoring functions or higher FDR%, however, these proteins are identified in all four of the biological replicates analyzed by AP-MS and therefore could be biologically relevant. Importantly, all the above proteins identified were part of the IP sample that was able to extend *T. brucei* telomeric repeats using synthetic TTAGGG as substrates in activity assays (**Figure 1F**), indicating that the proteins in this IP are part of an active telomerase complex.

### 3.4 The Unique C/D Box Domain in *T. brucei* telomerase RNA is bound by snoRNPs

Studies on vertebrate telomerase RNA have highlighted the importance of Cajal bodies as sites of telomerase maturation and assembly. Cajal Bodies are particularly linked with small nuclear ribonucleoprotein (snRNP) and small nucleolar ribonucleoprotein (snoRNP) biogenesis. Vertebrate telomerase RNAs contain a box H/ACA snoRNA-like domain at its 3’ end, which is bound by core snoRNPs of the H/ACA snoRNA family (Mitchell et al., 1999; Ghanim et al., 2021). The core H/ACA proteins bound to hTR are, dyskerin, NHP2, and GAR1. These H/ACA box binding proteins have been shown to be required for proper telomerase biogenesis and assembly *in vivo* (Egan and Collins, 2012). Mutations in these core H/ACA RNPs have been found to be associated with telomere shortening phenotypes and human disease (Kong et al., 2013).

In contrast to hTR, the telomerase RNA in *T. brucei* contains a unique C/D snoRNA-like domain (**Figure 4A** bottom). Core C/D box RNPs, like NOP58, have previously been shown to interact with *Tb*TR (Gupta et al., 2013). NOP58 is a 57 KDa protein that contains a coiled-coil (CC) domain and a NOP domain (**Figure 4A** top). NOP58 is highly conserved across eukaryotes and plays important roles in ribosomal RNA (rRNA) processing (Barth et al., 2008). In our AP-MS analysis of *Tb*TERT, we identified three core C/D box binding proteins: NOP58, NOP56, and Fibrillarin (NOP1) (**Table 2, Supplementary Table S1**). To validate the interaction with NOP58, we performed a co-immunoprecipitation (Co-IP) assay of *Tb*TERT and detected *Tb*TERT and NOP58 through western blotting (**Figure 4B** top). For further confirmation of this interaction, we performed Co-IPs of *Tb*TERT in both WT and ΔC/D box mutant cell lines. In the WT cells, NOP58 is present in the IP product, while in the ΔC/D mutant cells, NOP58 is absent **(Figure 4B** bottom). Since *Tb*TERT is present in both WT and ΔC/D IPs, this suggests that NOP58 does not have a direct interaction with *Tb*TERT, but binds with the RNA moiety (TR) of *T.brucei* telomerase. This data also supports the fact that the C/D box motif is required for the interaction between *Tb*TR and NOP58.

**Figure 4:**
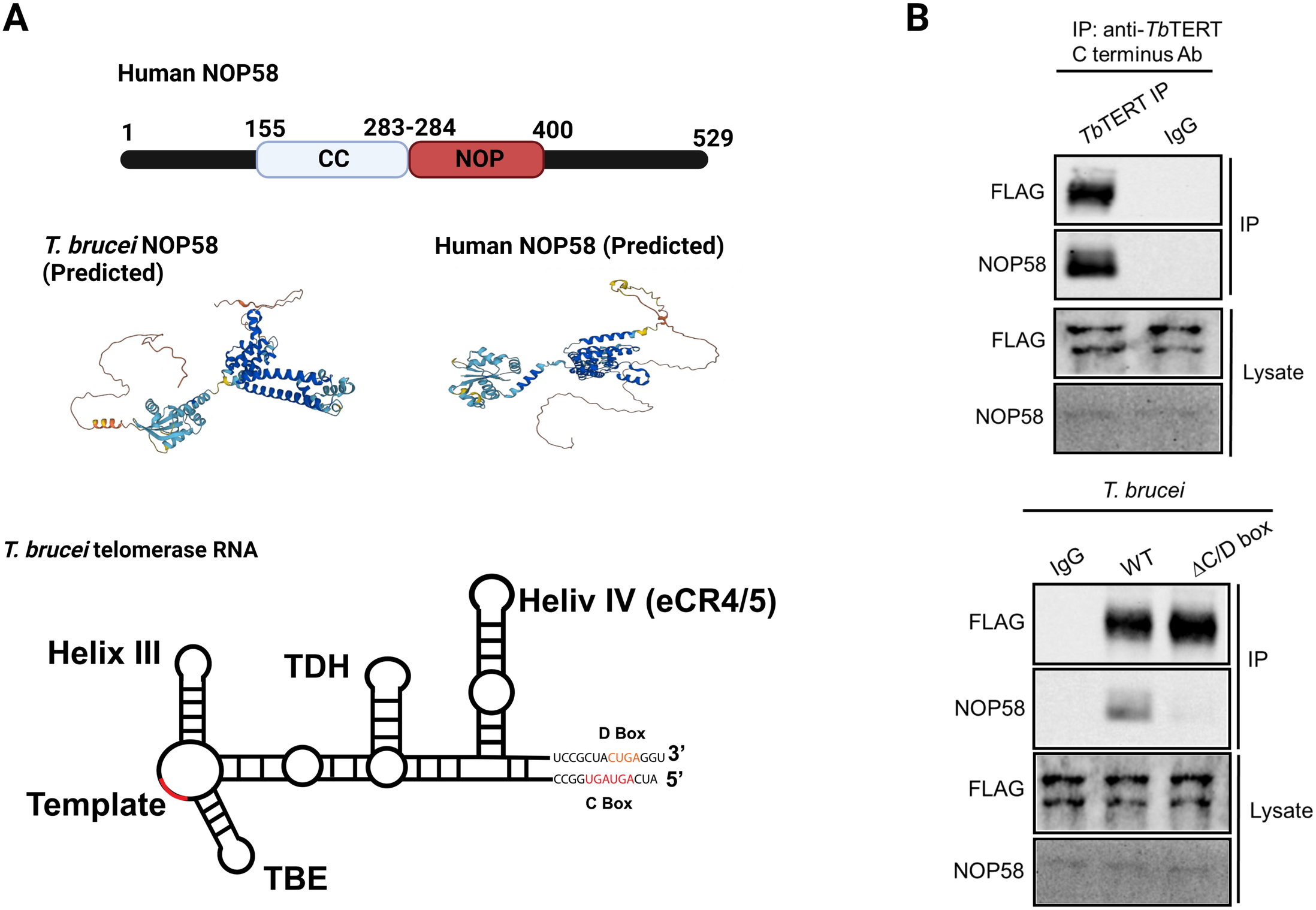
NOP58 interacts with the *T. brucei* telomerase complex. **(A)** Domain structure of human NOP58. Predicted secondary structure models for *T. brucei* and human NOP58 obtained from AlphaFold (Jumper et al., 2021). Dark blue represents a very high model confidence (pLDDT >90), light blue confident (90 >pLDDT >70), yellow low confidence (70 >pLDDT >50), orange very low confidence (pLDDT <50). Secondary structure model of *T. brucei* telomerase RNA. The C/D box binding motif is highlighted. **(B, top)** Co-IP assay using WT *T. brucei* cell lysate. IP antibody: anti- *Tb*TERT C terminus; Western blot antibodies: anti-FLAG and anti-NOP58. **(B, bottom)** Co-IP using both WT and ΔC/D box mutant cells. IP antibody: anti-*Tb*TERT C terminus; western blot antibodies: anti-FLAG and anti-NOP58.

**Table 2.** Core C/D snoRNPs identified in *Tb*TERT complex

| Protein | Accession number<br>(UniProtKB) | Spectral counts | Unique peptides |
| --- | --- | --- | --- |
| NOP58 | Q38F23 | 25 | 12 |
| NOP56 | Q580Z5 | 24 | 10 |
| Fibrillarin/<br>NOP1 | Q388E0 | 8 | 3 |

## 4. DISCUSSION

Eukaryotic microbes, such as *T. brucei*, rely on constitutive telomerase activity to sustain their proliferation in their hosts and to maintain the integrity of their subtelomeric virulence genes. Studying these interacting partners of telomerase in *T. brucei* is necessary to characterize the mechanism of telomerase mediated telomere maintenance in these parasites. Like any other RNA component of telomerase, *T. brucei* telomerase RNA in the holoenzyme acts as a structural scaffold that accessory proteins can bind to (Dey and Chakrabarti, 2018). Interacting proteins of telomerase in *T. brucei* have not been extensively identified or studied. We have utilized a mass spectrometry approach to identify and characterize endogenous interactions of *Tb*TERT in BF *T. brucei* cells. However, characterization of dynamic RNA-protein interactions like the one in telomerase complex comes with a challenge – that several of these interaction partners could be only transiently bound and therefore may not represent the complex interactions in its entirety. Nonetheless, this affinity purification based proteomic characterization of *T. brucei* telomerase RNP complex provides a global view of the cellular protein interactome landscape that can be used for in-depth functional characterization of telomerase complex proteins in parasites.

Some potential limitations of this study are the use of affinity purification methods. These methods are good at isolating strong interactions of our bait protein *Tb*TERT, but as mentioned above, weaker or more transient interactions may be missed by this analysis. Also, proteins of low abundance, such as TERT, could be difficult to enrich in affinity purified complexes even by targeted approaches like RNA-targeted APEX based proteomic approach recently employed for human telomerase (Han et al., 2020). For that reason, it is possible that several telomerase RNA and TERT associated proteins that are identified in this proteomic screen showed low level of enrichment in the IP complex, as evident from MS. Since IP experiments using MS provide a sensitive and accurate way of characterizing protein complexes, the quality of antibody may also play a role in isolating and analyzing specific interactions. The custom anti-*Tb*TERT polyclonal antibody used in the IP experiments was cross-reactive to other proteins (data not shown), however, the binding specificity to the endogenous bait protein *Tb*TERT was successfully confirmed using Co-IP and Western blot as *Tb*TERT was detected as a single, discrete band. Additionally, detection of the telomerase-associated proteins in all biological replicates added confidence to the current approach.

Apart from its telomeric functions in the nucleus, the telomerase catalytic subunit, TERT, was found to shuttle to other subcellular compartments, such as mitochondria, particularly in response to cellular stress. Accordingly, in addition to detection of nuclear telomerase-associated proteins, several mitochondrial proteins were highly enriched in IP samples, including *Tb*UMSBP2 (Klebanov-Akopyan et al., 2018). A protein involved in replication and segregation of the mitochondrial DNA, indicating potential mitochondrial function of *Tb*TERT in *T. brucei*. Significantly enriched GO terms from the analysis of the *Tb*TERT interactome included, telomere maintenance via telomerase, cell cycle control, and chaperone binding. Similar terms and protein interactors have been previously observed for *Saccharomyces cerevisiae* telomerase (Lin et al., 2015). Telomere maintenance via telomerase is consistent with *Tb*TERTs known role in extending telomeres (Dreesen et al., 2005). Enrichment of proteins involved in cell cycle control highlight potential factors involved in the cell cycle specific regulation of *Tb*TERT. Specifically, APC3 and CDC16 were significantly enriched in the *Tb*TERT interactome. APC3 and CDC16 are core components of the anaphase-promoting complex (APC), which is a 1.5 MDa ubiquitin ligase complex that regulates sister-chromatid separation and the cells exit from mitosis (Peters, 2006). In *S. cerevisiae*, the APC has been shown to degrade the telomerase recruitment subunit, Est1p to regulate telomere maintenance (Ferguson et al., 2013). Whether an analogous mechanism exists in *T. brucei* remains to be explored.

Chaperone proteins such as Dnaj, which is a major co-chaperone for HSP70 and HSP60 were also found to be significantly enriched in the *Tb*TERT interactome. Both Dnaj and HSP60 have previously been found to associate with human telomeres (Nittis et al., 2010). HSP60 is a predominately mitochondrial chaperone, where it works to maintain protein homeostasis (Caruso Bavisotto et al., 2020). In human cells, TERT has been previously reported to localize to the mitochondria and guard cells against oxidative stress (Ahmed et al., 2008). Human TERT has also been shown to associate with HSP60 and act independently of the TR in the mitochondria (Sharma et al., 2012). *Tb*TERT’s association with HSP60 suggests a pool of *T. brucei* telomerase may also be localized in the mitochondrion.

In addition to proteins involved in cell cycle control and chaperones, core C/D snoRNP proteins, NOP58, NOP56, and Fibrillarin (NOP1) were also identified in the *Tb*TERT interactome. Our co-IP western blot data validates the interaction of NOP58 with the *T. brucei* telomerase complex. NOP58 interacts with NOP56 and NOP1 to form a subcomplex, which participates in rRNA processing (Barth et al., 2008). Core C/D box binding proteins, like NOP58, have been previously shown to interact with the *Tb*TR (Gupta et al., 2013). Our study supports these findings and shows that NOP58 specifically interacts with the C/D box motif in the *Tb*TR. Our data also indicates that NOP58 binds with *Tb*TR only and does not have a direct interaction with *Tb*TERT. The C/D box motif in *Tb*TR is unique and lacking in higher eukaryotes. The TR in *Leishmania* also contains a C/D box motif (Vasconcelos et al., 2014). The conservation of the C/D box motif in the TR of these parasites could indicate a novel mechanism for telomerase biogenesis and processing, mediated by C/D box binding proteins, in these kinetoplastid parasites.

The work described here provides the first analysis of the *Tb*TERT interactome. We have identified previously known and novel interactors of *Tb*TERT. We were able to confirm NOP58s interaction with the *T. brucei* telomerase complex, which supports earlier studies (Gupta el al., 2013). Taken together our study lays the foundation for future studies into the mechanism of telomerase mediated telomere maintenance in *T. brucei*. Future improvements are needed to develop a telomerase RNA - tagged proteomic mapping approach in *T. brucei* to validate endogenous interactions identified by this method and also detect new RNA-specific interactions. Future studies should also benefit from investigating interactomes from other *T. brucei* developmental stages since it appears from our recent study that *T. brucei* telomerase function is developmentally regulated (Dey et al., 2021). Therefore, characterizing stage-specific interactomes can provide novel insights into regulatory mechanisms that can affect rate of proliferation and telomerase activity in *T. brucei*.

## Author Contributions

JD, BL and KC designed experiments. JD, AR, N and AS conducted experiments. SX, AT, BL, and KC contributed critical reagents and suggestions. JD, AR, N and DW analyzed data. JD and KC wrote the manuscript. All authors revised and approved the final manuscript.

## Acknowledgments

We thank Chakrabarti lab members at UNC Charlotte and Li lab members at the Cleveland State University for their help and support with this work and Dr. Jun-tao Guo from the Department of Bioinformatics and Genomics at UNC Charlotte for help with structure-based prediction analysis using PDBeFold and DaliLite. Figures were created using BioRender.com

## Supplementary Material

The supplementary material for this article can be found online.

## Data Availability Statement

The datasets presented in this study can be found in online repositories. The names of the repository/repositories and accession number(s) can be found at: https://www.ebi.ac.uk/pride/archive/, PXD038235.

