## Supplemental figure 1 for "Proteomic analysis defines the interactome of telomerase in the protozoan parasite, *Trypanosoma brucei*"

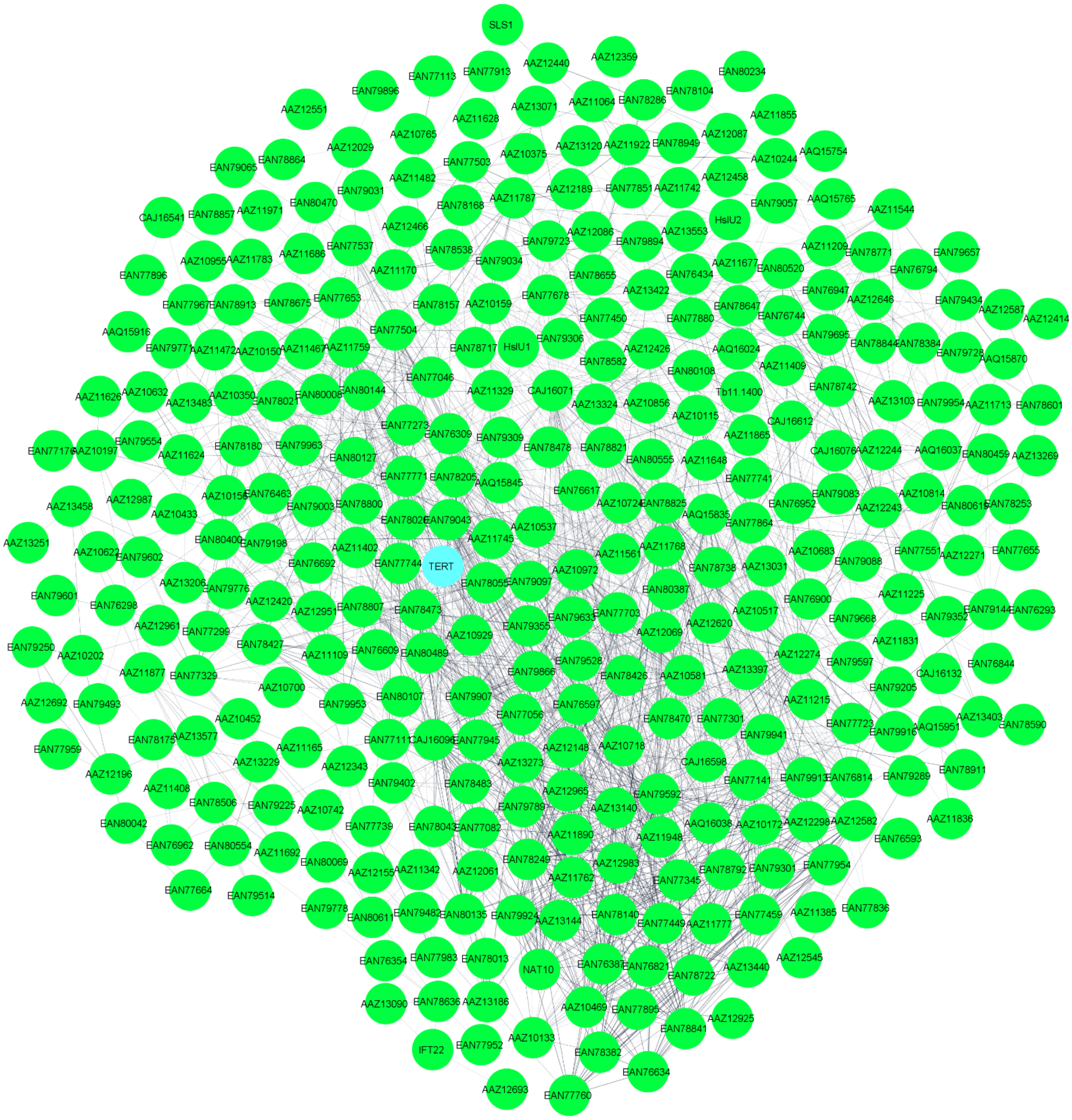


**Supplementary Figure S1:** Protein-protein interaction map of *Tb*TERT interactome. Protein-protein interaction network was generated using STRING and visualized by cystoscope. Line thickness denotes the strength of the interaction (confidence PPI, threshold: 0.4, medium confidence). *Tb*TERT protein is highlighted in blue.
