## Supplemental Table 1 for "Proteomic analysis defines the interactome of telomerase in the protozoan parasite, *Trypanosoma brucei*"

**Supplementary Table 1. *T. brucei* TERT interacting partners – known homologs**

| Gene/Protein | Gene ID | UniProt ID | Unique Peptides | *DaliLite Z-score | minusLog10(P-value) /log2 Fold Change | |
|---|---|---|---|---|---|---|
| | | | | | TbTERT Ab IP-MS | FLAG IP-MS |
| Heat shock protein 90 (**HSP90**), putative | Tb927.3.3580 | Q57W94 | 35 | 31.0 | 1.52/ -0.64 | .12/ -0.72 |
| AAA family ATPase, **CDC48** homolog of *S.cerevisiae*) | Tb927.10.5770 | Q38B27 | 33 | 49.4 | 0.226/ 0.113 | 0.14/ -0.61 |
| RuvB-like DNA helicase, (**PONTIN**) putative | Tb927.4.1270 | Q581V4 | 17 | 41.9 | 0.136/ -0.09 | 0.73/ -2.3 |
| RuvB-like DNA helicase, (**REPTIN**) putative | Tb927.4.2000 | Q583J3 | 13 | 39.4 | 1.58/ -0.82 | 0.02/ -0.04 |
| Nucleolar protein 58 (**NOP58**), putative | Tb927.9.5320 | Q38F23 | 12 | 26 | 0.27/ 0.75 | 1.08/ 2.25 |
| Poly(A)-specific ribonuclease **PARN**, putative | Tb927.9.13510 | Q38D76 | 11 | 32.8 | 0.87/ 1.3 | 0.65/ 2.2 |
| Replication factor A protein 1 | Tb927.11.9130 | Q384B5 | 10 | 20.8 | 0.11/ -0.18 | 0.34/ 0.56 |
| Nucleolar protein 56 (**NOP56**), putative | Tb927.8.3750 | Q580Z5 | 10 | 27.2 | 0.4/ -0.15 | 0.8/ 1.90 |
| Telomerase Reverse Transcriptase (***Tb*TERT**) | Tb927.11.10190 | Q383R0 | 5 | 22.9 | 3.0/ 1.98 | 0.38/ 0.49 |
| La protein (**p65**) homolog, putative | Tb927.10.2370 | Q38C07 | 3 | 8.6 | 0.19/ -0.63 | 0.09/ -0.49 |
| **Fibrillarin**, putative | Tb927.10.14630 | Q388C9 | 3 | 33.5 | 0.26/ 0.27 | 0.78/ 1.97 |

*'Significant similarities' have a Z-score above 2; they usually correspond to similar folds when searched against entire PDB matches.
